# In-depth Temporal Transcriptome Profiling of Monkeypox and Host Cells using Nanopore Sequencing

**DOI:** 10.1101/2022.11.28.518226

**Authors:** Balázs Kakuk, Ákos Dörmő, Zsolt Csabai, Gábor Kemenesi, Jiří Holoubek, Daniel Růžek, István Prazsák, Virág Éva Dani, Béla Dénes, Gábor Torma, Ferenc Jakab, Gábor E. Tóth, Fanni V. Földes, Brigitta Zana, Zsófia Lanszki, Ákos Harangozó, Ádám Fülöp, Gábor Gulyás, Máté Mizik, András Attila Kiss, Dóra Tombácz, Zsolt Boldogkői

## Abstract

The recent Monkeypox outbreak showed the importance of studying the basic biology of orthopoxviruses. However, the transcriptome of its causative agent has not been investigated before neither with short-, nor with long-read sequencing approaches. This Oxford Nanopore long-read RNA-Sequencing dataset fills this gap. Our direct cDNA and native RNA sequencing data enable the in-depth characterization of the transcriptomic architecture and dynamics of the gene expressions of monkeypox virus; and also the deeper understanding of the changes it causes in the host cells on a transcriptome level.

## Background & Summary

Monkeypox virus (MPXV) belongs to the *Poxviridae* family, which contains many viruses that infect various animal taxa including invertebrates, reptiles, and mammals. MPXV is the member of the human pathogenic *Orthopoxvirus* genus, which also includes the cowpox virus, the vaccinia virus (VACV) and the highly dangerous variola virus, the causative agent of smallpox^1,2^. Smallpox infections caused millions of deaths throughout the history until a global vaccination program has successfully eradicated the virus from the human population^3^. Infections of MPXV, have also been reported, although with lower mortality and milder morbidity^3^. Human MPXV infections were localized to Central and West Africa during the last decades, except for some rare cases. However, due to a recent outbreak, a growing number of cases were reported from countries where the disease is not endemic^4–6^. The genomic monitoring of the 2022 MPXV outbreak has demonstrated that the recently circulating MPXV strain is probably related to the less likely pathogenic West African clade of MPXVs and forms a highly divergent new clade with an elevated mutation rate^7–9^.

The orthopoxviruses are one of the largest of all animal viruses with their sizeable, brick-shaped, and membrane-coated virions of about 200-300 nm in diameter and their large, linear double-stranded DNA genome, with around 200 kilobase pairs in size^10^. In contrast to most mammalian DNA viruses (such as herpesviruses and adenoviruses), which replicate in the nucleus, poxviruses remain in the cytoplasm. Viral replication and transcription of MPXV genes take place within compartments called “viral factories” independently of the host cell^11^. This peculiar feature draws attention to how MPXV regulates the gene expression of the host cell.

The transcriptional effect of MPXV infection on different cell types has been characterized using micro-array-based techniques^12–14^. Rubins and colleagues used a high-resolution poxvirus-human microarray covering 24h of infection and classified all MPXV genes for the first time according to their temporal expression^15^. They also compared the expression profile of MPXV to VACV and found that only the minority of transcripts are species-specific^15^. And though recent studies have re-evaluated these data using comparative pathway analyses, the detailed transcriptomic characteristics of MPXV-infected cells remains undescribed^16^. Thus, while micro-array-based techniques reveal useful insights, they are unable to resolve many aspects of the transcriptome, including the detection of the plethora of different transcript isoforms, which have been detected in closely related viruses, for example in VACV^17^.

RNA Sequencing has become the most widely applied method in transcriptome research. Short-read sequencing (SRS) techniques generate sufficient depth of sequencing and have a high accuracy, but transcriptome annotations may remain incomplete because of the fragmented nature of the sequenced cDNAs^18–20^. This is especially true in the case of viruses, which have gene-dense genomic regions where transcripts substantially overlap each other. Additionally, SRS has a severe limitation for distinguish the different transcript isoforms^21^. Long-read sequencing methods (LRS), including Pacific Biosciences and Oxford Nanopore Technologies (ONT) offer an alternative for transcriptome sequencing that enables the recovery of full-length RNA molecules, which is crucial for a precise transcriptome annotation^22^. Although these methods generate less reads and have relatively higher error rates compared to SRS, with sufficient coverage, the transcriptome annotation of well-annotated genomes, like MPXV becomes possible^23–26^. Moreover, ONT can sequence native RNAs directly (dRNA-seq), hence it avoids the generation of false products of the reversetranscription or PCR steps during the library preparation. A drawback of dRNA ONT sequencing technique, however, its inability to detect 5’ termini of mRNAs^27^. This problem can be circumvented with the combined usage of 5’-end sensitive PCR-free direct cDNA sequencing methods dcDNA)^23,28–30^. Moreover, direct cDNA-seq can be used to accurately quantify gene expression, as it is not affected by biases introduced in the RT-PCR of traditional PCR-cDNA-sequencing^31^.

As of now, the transcriptome of only a few poxvirus has been analyzed by next generation sequencing methods, including the VACV a model for orthopoxviruses and a close relative of MPXV^32–36^. LRS methods have been used to redefine the highly intricate structure of VACV transcriptome^37^, moreover the dynamic gene expression changes were analyzed in detail during the time course of infection^17,38,39^. However, to our knowledge there is a lack of high-throughput RNA sequencing studies regarding the MPXV transcriptome. Hence, our goal in this work is to present an LRS dataset for an accurate transcriptome annotation of MPXV.

In this study, the transcriptomes of the MPXV along with its host cell were sequenced using an Oxford Nanopore Technologies (ONT) MinION long-read sequencing device. Two sequencing approaches were utilized in this study: a dcDNA-seq of 6 different time-points (1-, 2-, 4-, 6-, 12- and 24-hours post infection) from the virus-infected cells, each with 3 biological replicates, and a dRNA-seq library from a mixture of the time-point samples.

This dataset can be used for the analysis of temporal transcriptomes of MPXV and the infected cells. Since even short-read transcriptomic data are completely missing of MPXV, our long-read RNA-seq dataset should serve as a gap-filler and will enable the in-depth characterization of its transcriptome.

## Methods

**Figure 1** shows the detailed workflow of the study.

**Figure 1.**
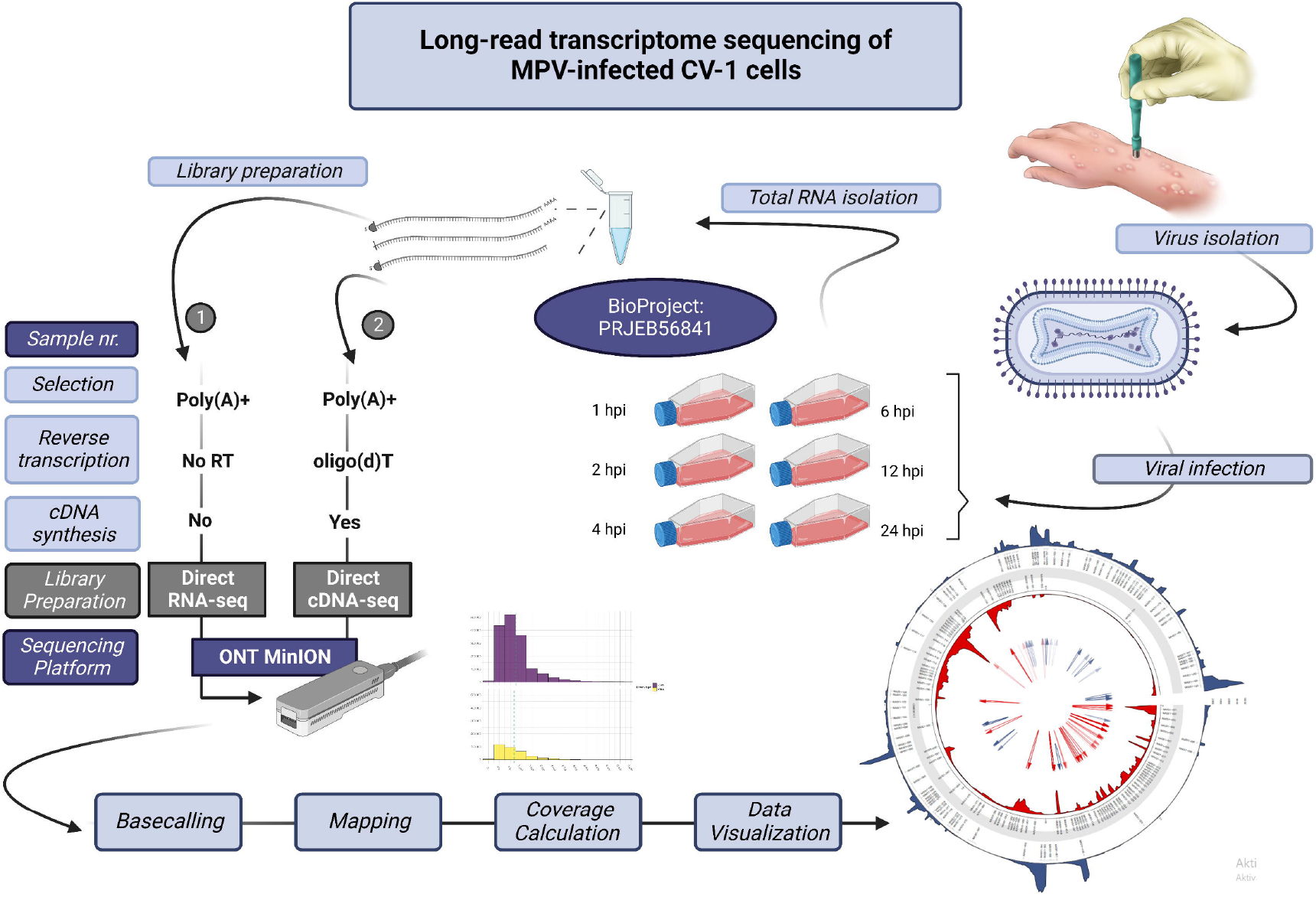
General overview of the study. The general overview of the study is represented in this figure, which was created with Biorender (BioRender.com).

### Cells

CV-1 (CCL-70, African green monkey, kidney) cell line was used which was obtained from American Type Culture Collection (ATCC). For the experiment 75 cm^2^ tissue culture flasks (CELLSTAR®; Greiner Bio-One GmbH, Frickenhausen, Germany) were plated with 2 x 10^5^ cells in Minimum Essential Medium Eagle culture medium (MEM) with 10% fetal bovine serum (FBS). The CV-1 cells were cultivated until ~80% (~1.2 x 10^6^) confluency at 37°C in humified 5% CO_2_ atmosphere. Before the infection, the monolayer was washed with 1 X PBS (Thermo Fisher Scientific, Waltham, MA, USA).

### Collection, detection, isolation and propagation of the virus

The MPXV (MPXV_NRL 4279/2022) was isolated from skin lesions and kindly provided by Dr. Jirincova (The National Institute of Public Health, Prague, Czech Republic). All procedures with infectious materials were performed under BSL-4 conditions at the National Laboratory of Virology, University of Pécs. The virus was passaged once on CV-1 cells to reach a sufficient amount of infective particles. The same batch of working stock was used during the experiment. The viral titer of the working stock was determined with plaque assay on CV-1 cells. Non-infected control cultures were inoculated with MEM and treated the same way as the infected ones. For the infection, 2 ml MPXV with 5 plaque-forming units (pfu)/cell (MOI = 5) was used, which was diluted with MEM to reach the sufficient concentration. Cells were incubated with monkeypox inoculum at 37°C for 1 hour while were shaken gently in every ten minutes. The virus inoculum was removed, then the cell monolayer was washed once with 1 x PBS. For the flasks 10 mL MEM medium was added which was supplemented with 2% FBS, 2 mM L-glutamine and 1% penicillin and streptomycin solution. The cells were incubated at 37°C for 1, 2, 4, 6, 12 and 24 hours in a humidified 5% CO_2_ atmosphere. Each time, the experiment was done in triplicate and subjected to direct cDNA RNA sequencing. Prior to direct RNA sequencing extra flask was used to sample the following time points: 2-, 6-, 12- and 24-hours post-infection. Direct RNA sequencing samples were treated without replicates. After the incubation, the supernatant was removed, and the cells were washed with PBS. The dry flasks were stored at −80°C until further processes. The cells were washed and scraped down into lysis buffer and transferred to 1,5 mL Eppendorf Tubes^®^ (Thermo Fisher Scientific, Inc.).

### Isolation of total RNA

Total RNA was purified from the MPXV-infected and from mock-infected CV-1 cells at various time points after infection from 1 to 24 hours. For this, the NucleoSpin RNA Kit (Macherey-Nagel) was used, following the manufacturer’s recommendations. Briefly, cells were collected by centrifugation (1000 x g), then 350μl RA1 lysis buffer (part of the NucleoSpin RNA Kit) and 3.5μl β-Mercapthoethanol (Sigma Aldrich) were added to the samples and then, mixtures were centrifuged at 11,000 x g for 1 min in NucleoSpin Filter tubes. Filters were discarded, and the lysate was washed using 70% EtOH (350μl) on NucleoSpin RNA Column with centrifugation at 11,000 x g for 30sec. Membrane Desalting Buffer (350μl, from the NucleoSpin RNA Kit) was then added to desalt the membrane, which was finally dried with centrifugation (11,000 x g). Residual DNA was removed using rDNase enzyme [rDNase:rDNase reaction buffer (1:9 ratio, NucleoSpin Kit)]. The enzymatic reaction was carried out at room temperature (RT) for 15min. The NucleoSpin Kit’s RAW2 Buffer (200μl) was used on the NucleoSpin Filter, which inactivated the enzyme. After a short centrifugation (11,000 x g, 30min) the Filter was placed in a new Eppendorf tube. The next washing step was carried out with RAW3 Buffer (600μl, from the NucleoSpin RNA Kit) and centrifugation (11,000 x g, 30min). This step was repeated with 250μl RAW3 Buffer. The purified total RNA samples were eluted from the Filter in 60μl nuclease-free water (NucleoSpin RNA Kit) and they were stored at −80°C (**Table 1**).

**Table 1.**
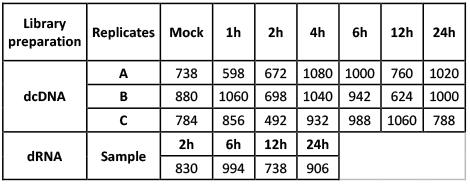
RNA quantities. Obtained yield of total RNA samples (in ng/μl). Upper panel: RNA samples used for dcDNA sequencing; lower panel: RNAs from which the dRNA-seq sample was mixed.

### Poly(A) selection

Polyadenylated RNA was enriched using the Lexogen’s Poly(A) RNA Selection Kit V1.5. This method is based on oligo(dT) beads, which hybridize RNAs with polyadenylated 3′ ends, but RNAs without poly(A) stretches (e.g. rRNAs) do not captured by the beads and therefore, they will be washed out. The applied protocol is as follows: the beads (from of the Lexogen Kit) were resuspended and 4μl for each RNA samples was used. Beads were collected in a magnet, and the supernatant was discarded. RNAs were resuspended in Bead Wash Buffer (75μl, Lexogen Kit) and then were placed on the magnet, and supernatant was discarded. This washing step was repeated. Beads were resuspended in RNA Hybridization Buffer (20μl, Lexogen Kit). Ten μg from the total RNA samples were diluted to 20μl in nuclease-free water (UltraPure™, Invitrogen) and then they were denatured at 60°C for 1min. Denatured RNA samples were mixed with 20μl beads. The mixtures were incubated in a shaker incubator with 1250 rpm agitation at 25°C for 20min. Next, the samples were placed in a magnetic rack. Supernatant was discarded, the tubes were removed from the magnet, the collected samples were resuspended in 100μl Bead Wash Buffer (Lexogen Kit), and finally, they were incubated for 5min at 25°C with 1250 rpm agitation. Supernatant was discarded and this washing step was repeated once. Beads were resuspended in 12μl nuclease-free water, then kept at 70°C for 1min. After this incubation step, tubes were placed on a magnetic rack and supernatant, containing the polyadenylated fraction of RNA samples were placed to new DNA LoBind (Eppendorf) tubes (**Table 2**). Samples were stored at −80°C.

**Table 2.** Amount RNA samples (in ng/μl) obtained after PolyA purification. Upper panel: RNA samples used for dcDNA sequencing; lower panel: Polyadenylated RNAs for dRNA sequencing.

| Library preparation | Replicates | Mock | 1h | 2h | 4h | 6h | 12h | 24h |
| --- | --- | --- | --- | --- | --- | --- | --- | --- |
| dcDNA | A | 12,8 | 14,7 | 14,2 | 11,7 | 12,3 | 13,1 | 11,8 |
|  | B | 11,8 | 11,4 | 13,1 | 11,6 | 12,9 | 14 | 13,5 |
|  | C | 12 | 13,4 | 14,9 | 11,2 | 12,5 | 11,4 | 14,3 |
| dRNA | Sample | 2h | 6h | 12h | 24h |  |  |  |
|  |  | 12,4 | 12,2 | 14,4 | 12,4 |  |  |  |

### Direct cDNA sequencing

Direct (d)cDNA libraries were generated with the aim of analyzing the dynamic pattern of MPXV transcripts and the effect of viral infection on the host cell gene expression profile. RNA samples from different time points (1, 2, 4, 6, 12 and 24h p.i., and from the mock, three biological replicates from each) were used individually for library preparation. The ONT’s Direct cDNA Sequencing Kit (SQK-DCS109, ONT) was applied according to the manufacturer’s recommendations. Briefly, first-strand cDNAs were synthesized from the polyA(+) RNA samples using the Maxima H Minus Reverse Transcriptase enzyme (Thermo Fisher Scientific) and the SSP and VN primers (supplied in the ONT kit). The potential RNA contamination was eliminated by applying RNase Cocktail Enzyme Mix (Thermo Fisher Scientific).

The second cDNA strands were generated with LongAmp Taq Master Mix (New England Biolabs). The ends of the double-stranded cDNAs were repaired with NEBNext End repair /dA-tailing Module (New England Biolabs) and then the adapters were ligated using the NEB Blunt/TA Ligase Master Mix (New England Biolabs). The Native Barcoding (12) Kit (ONT) was for multiplex sequencing **(Table 3)**. The samples (200 fmol/flow cell) were loaded onto MinION R9.4 SpotON Flow Cells (ONT, **Table 4**).

**Table 3.**
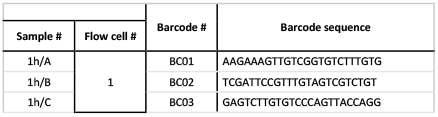
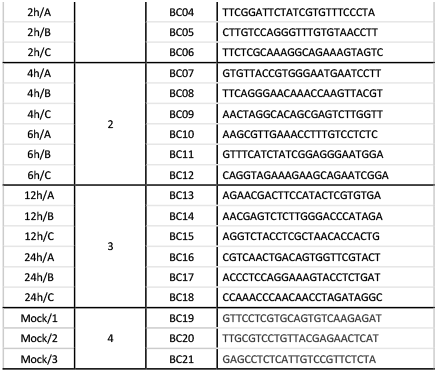
Sequencing barcodes used.

**Table 4.** Amount of libraries (in μl) that were used for sequencing. Two-hundred fmol dcDNA library mixture was loaded onto each of the Flow Cells (33.34 fmol/sample from viral infected samples and 66.67 fmol from the mock-infected libraries).

|  | Flow Cell # 1 |  | Flow Cell # 2 |  | Flow Cell # 3 |  | Flow Cell # 4 |
| --- | --- | --- | --- | --- | --- | --- | --- |
| $\mu\text{l}$ | 1h | 2h | 4h | 6h | 12h | 24h | Mock |
| A | 7,2 | 8,48 | 8,87 | 7,7 | 5,15 | 5,36 | 15,39 |
| B | 7,74 | 6,26 | 8,81 | 10,71 | 6,87 | 6,32 | 15,85 |
| C | 7,44 | 5,53 | 7,09 | 7,65 | 5,6 | 4,56 | 13,66 |

### Direct RNA sequencing

Direct RNA sequencing (SQK-RNA002; Version: DRS_9080_v2_revO_14Aug2019, Last update: 10/06/2021) was used to sequence the native RNA strands to avoid any potential bias from reverse transcription or PCR. Fifty ng (in 9 μl) from a mixture of polyA(+) RNAs from various time points (2, 6, 12 and 24h p.i.) was used for library preparation. As a first step, 1 μl RT Adapter (110nM; ONT Kit) was ligated to the RNA sample using 3μl NEBNext Quick Ligation Reaction Buffer (New England BioLabs), 0.5μl RNA CS (ONT Kit), and 1.5μl T4 DNA Ligase (2M U/ml New England BioLabs) at RT for 10 min. The first cDNA strand was generated using SuperScript III Reverse Transcriptase (Life Technologies), as recommended by the Direct RNA sequencing (DRS) manual (ONT). The reaction was carried out at 50°C for 50 min and it was followed by the inactivation step at 70°C for 10 min. Next, the sequencing adapters (ONT’s DRS kit) were ligated to the cDNA at RT for 10 min using the T4 DNA ligase enzyme and NEBNext Quick Ligation Reaction Buffer. The dRNA library was sequenced on an R9.4 SpotON Flow Cell.

RNAClean XP beads and AMPure XP beads (both from Beckman Coulter) were used after each of the enzymatic reactions for washing the dRNA-seq and dcDNA-seq libraries, respectively.

### Bioinformatics

The generated sequencing reads were basecalled with the Guppy software (available at ONT’s community site https://community.nanoporetech.com/), with the following parameters: -*flowcell FLO-MIN106 --kit SQK-DCS109 --barcode_kits EXP-NBD114 --min_qscore 8 --recursive --calib_detect*. Based on a quality threshold of 8, the basecalled reads were separated into a ‘pass’ and a ‘fail’ group – the subsequent analyses were carried out on the *passed* reads. The *.fastq* files containing the *passed* reads for the respective samples were merged.

The resulting sequences were then mapped to a combined reference, containing the host genome (accession number: GCF_015252025.1) and the viral genome (ON563414.3), using *minimap2*^40^. The reference genomes were downloaded from the GenBank and ENSEMBL, respectively. The mapping parameters were the following: *minimap2-ax splice -Y -C5 --cs -MD -un -G 10000*.

The subsequent analyses were carried out within the R environment – all scripts are available in the GitHub repository (https://github.com/Balays/MPOX_ONT_RNASeq). The workflow implements functions from the tidyverse^41^ collection of R packages. The complete workflow can be re-run to produce all the analysis results, including generation of figures and tables. The first step in the MPOX-wf is to import the .bam files into the R workspace using Rsamtools^42^. Raw alignment counts were calculated using *idxstats*. Then reads with secondary alignments were filtered out, as these are putatively chimeric RNAs. Viral and host read counts, according to the mapping results (**Figure 2**) and read lengths (**Figure 3** and **Supplementary Figure S1**) were visualized with the ggplot2 package^43^. Next, per-base coverage values and their mean along a 100 nt window were calculated. The mean coverage on monkeypox genome in the dRNA sample and in the dcDNA samples (after *log10* normalization) was visualized using the circlize package^44^ (**Figure 4** and **Figure 5**, respectively). The links in the center of the circle represent transcripts, as in the connections of the 5’- and 3’-ends of the reads. These potential ‘transcripts’ were filtered to read count threshold of 10. The transparency of the links is correlated with the abundance of the ‘transcripts’.

**Figure 2.**
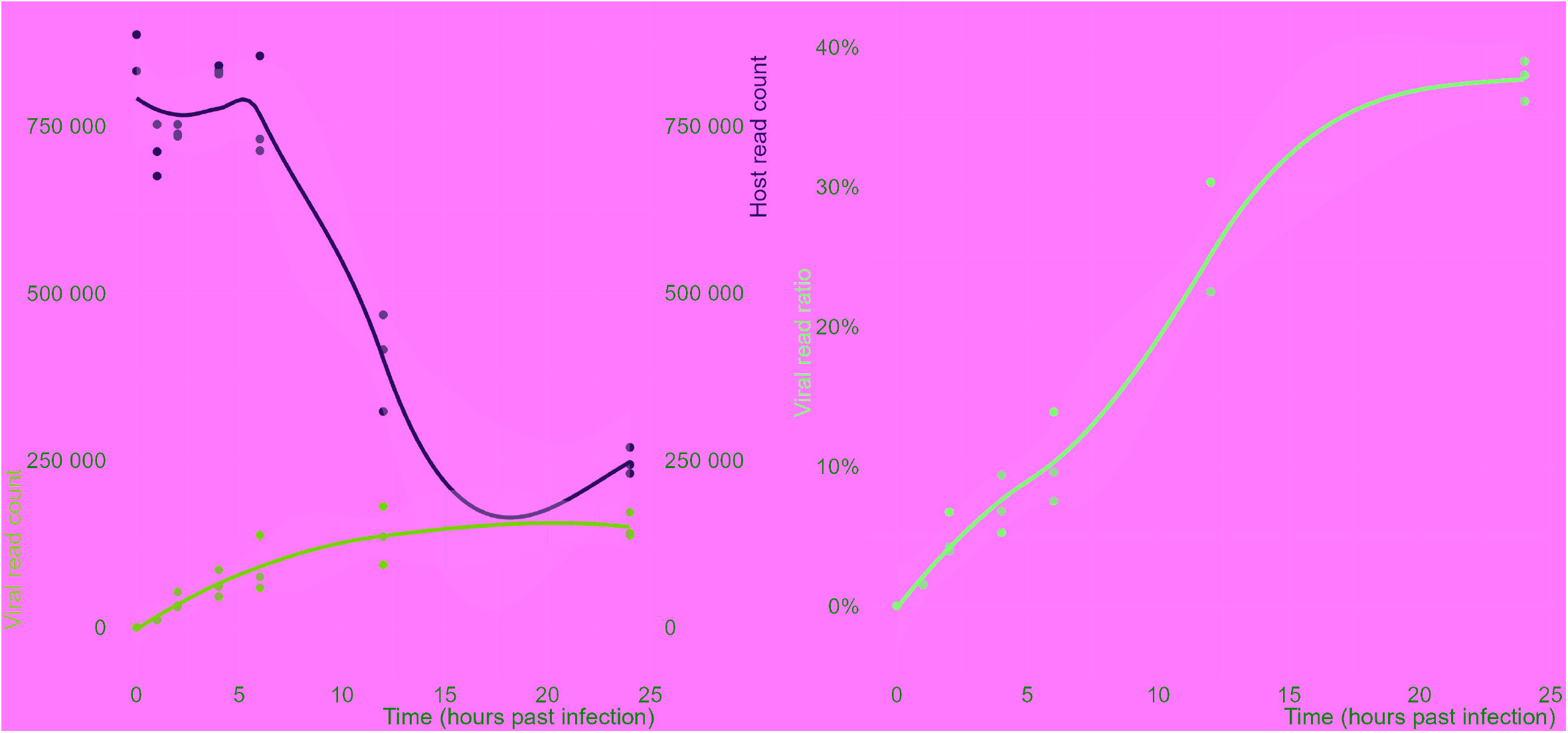
Sequencing read counts and viral read ratios.

**Figure 3.**
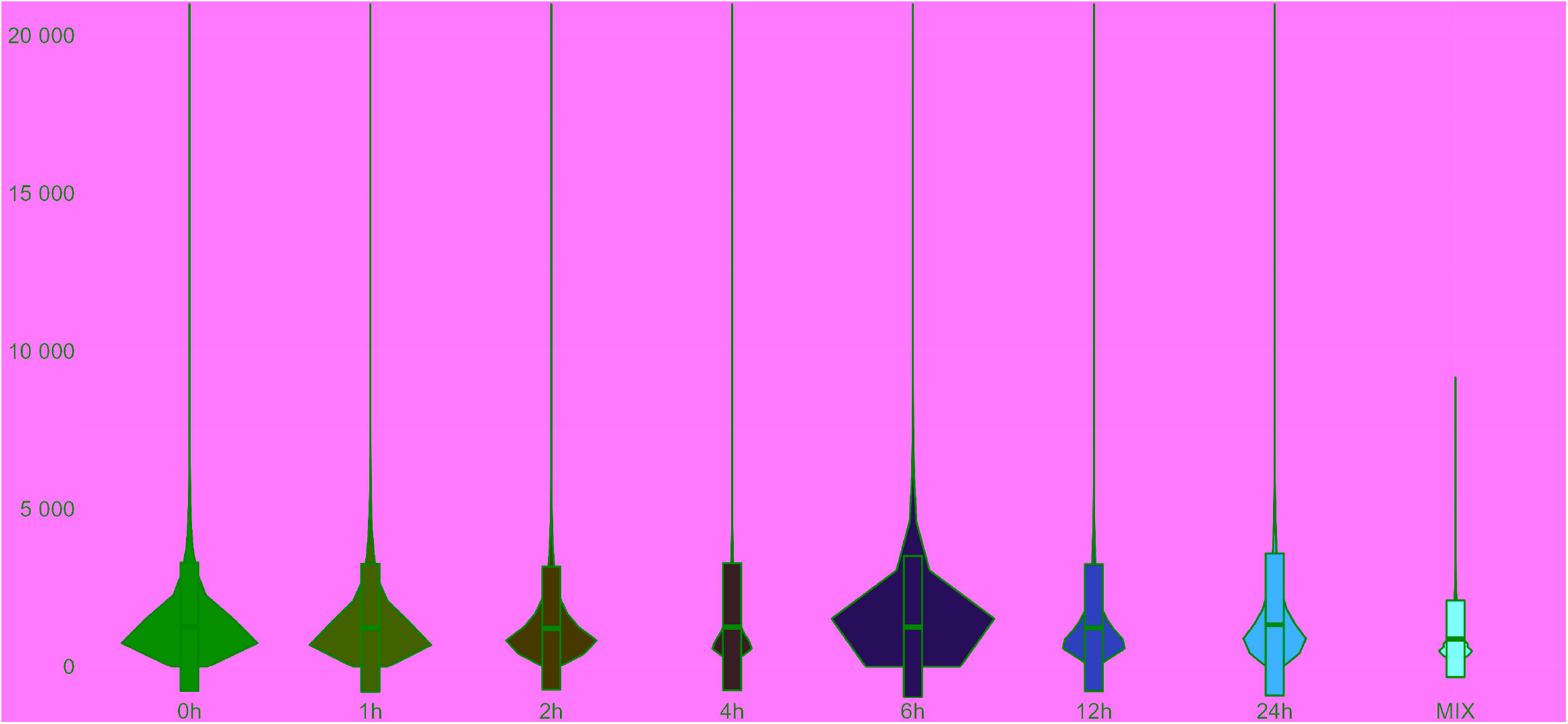
Read length distribution in the cDNA and the dRNA sequencing libraries.

**Figure 4.**
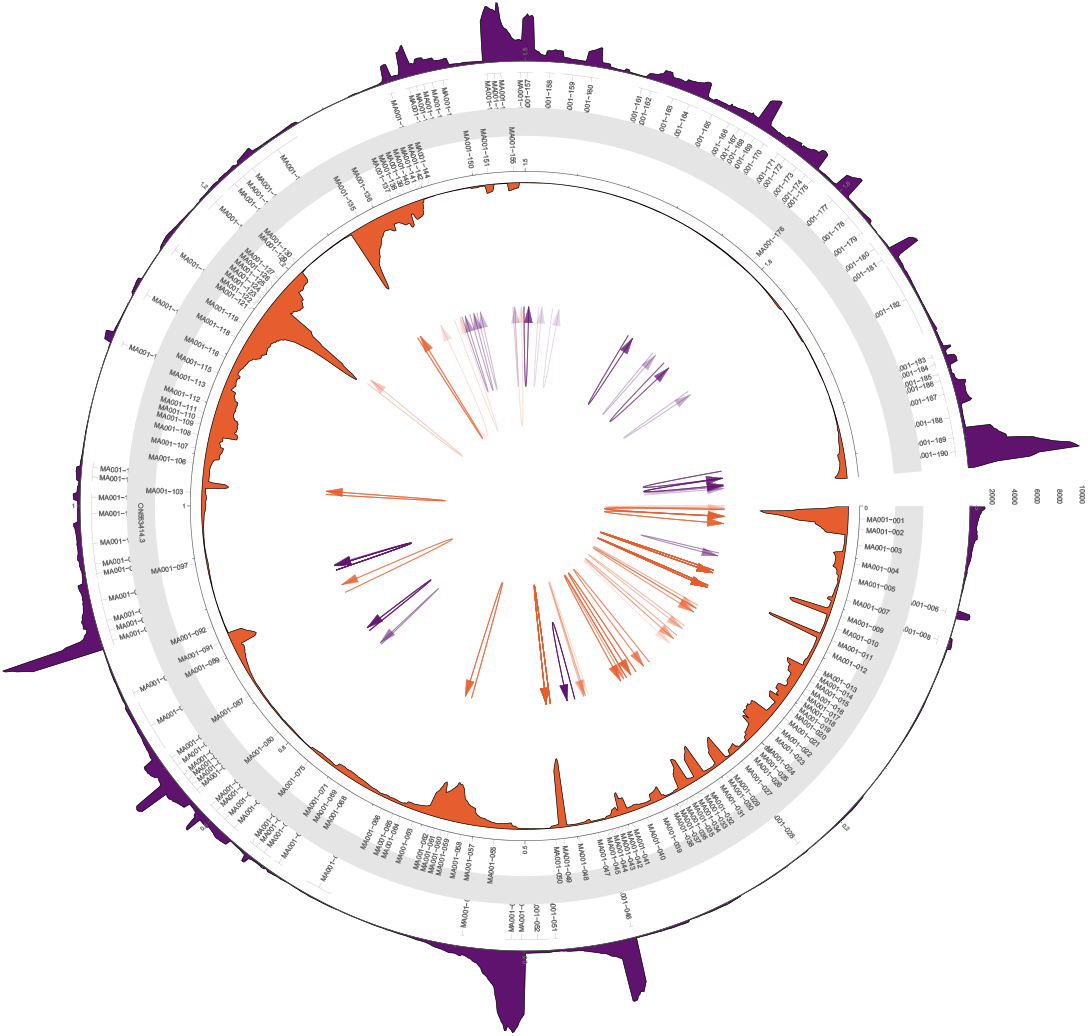
Coverage of the viral genome in the dRNA sequencing library. The mean coverage on monkeypox genome was calculated in a 100-nt window. The links in the center of the circle represent transcripts, as in the connections between the 5’- and 3’-ends of the reads. These potential ‘transcripts’ were filtered to read count threshold of 10. The transparency of the links is correlated with the abundance of the ‘transcripts’.

**Figure 5.**
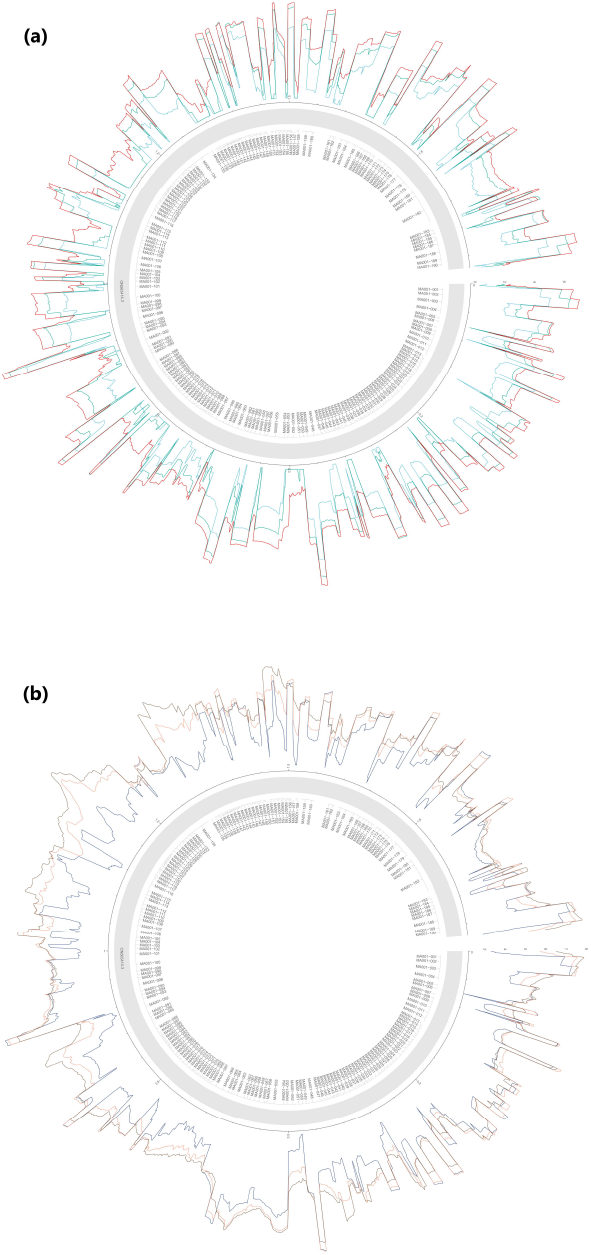
Coverage (normalized to *log10*) of the viral genome in the dcDNA sequencing library. (a) Coverage in 1-, 2- and 4-hours post-infection; **(b)** coverage in 6-, 12- and 24-hours postinfection.

## Data Records

The 21 dcDNA sequencing yielded a substantial amount of 16,150,173 reads that passed guppy’s QC filtering threshold of 8 (**Table 5**). These were mapped onto the combined host and viral reference (**Figure 2**, left panel). The distribution of read lengths is shown in **Figure 3** and the viral reads in **Supplementary Figure S1**. The mean of the read lengths did not change significantly, most of the reads were in the 800-1000 nt bin.

The ratio of viral reads showed a steady increase from around 1.5%±0.5% in the 1 hpi samples to 37% ±0.5% in the 24 hpi samples (**Figure 2**, right panel). The median coverage across the whole viral genome also increased: from 11 to 571 (**Figure 5**). The total read count peaked at 4- and 6-hours post-infection and decreased afterwards.

**Table 5.** Sequencing summary. The ‘Total read count’ column shows the number of QC pass reads. The ‘Host read count’ column shows the number of reads mapped onto the Vero genome; while the ‘Viral read count’ column shows the number of reads that were mapped to the ON563414.3 genome, and that did not have secondary alignments (as these are potentially chimeric reads). The ‘Viral read ratio’ column corresponds to the ratio of these non-chimeric viral reads and the ‘Total read count’ column.

| sample | hpi | Time | Mapped read count | Unmapped read count | Viral read count<br>nochimaera | Total read count | Host read count | Viral read ratio |
| --- | --- | --- | --- | --- | --- | --- | --- | --- |
| mock_A | 0h | 0 | 755484 | 60974 | 0 | 816458 | 0 | 0 |
| mock_B | 0h | 0 | 832329 | 70647 | 1 | 902976 | 832328 | 0 |
| mock_C | 0h | 0 | 886821 | 74969 | 1 | 961790 | 886820 | 0 |
| 1h_A | 1h | 1 | 685835 | 51024 | 10713 | 736859 | 675122 | 0,0156 |
| 1h_B | 1h | 1 | 763926 | 86085 | 11501 | 850011 | 752425 | 0,0151 |
| 1h_C | 1h | 1 | 722542 | 47980 | 10798 | 770522 | 711744 | 0,0149 |
| 2h_A | 2h | 2 | 770363 | 77972 | 32320 | 848335 | 738043 | 0,042 |
| 2h_B | 2h | 2 | 787439 | 46728 | 52820 | 834167 | 734619 | 0,0671 |
| 2h_C | 2h | 2 | 782981 | 54857 | 30664 | 837838 | 752317 | 0,0392 |
| 4h_A | 4h | 4 | 873504 | 40219 | 45877 | 913723 | 827627 | 0,0525 |
| 4h_B | 4h | 4 | 901540 | 77305 | 61162 | 978845 | 840378 | 0,0678 |
| 4h_C | 4h | 4 | 918908 | 47813 | 85903 | 966721 | 833005 | 0,0935 |
| 6h_A | 6h | 6 | 789700 | 36454 | 59224 | 826154 | 730476 | 0,075 |
| 6h_B | 6h | 6 | 992717 | 56782 | 137766 | 1049499 | 854951 | 0,1388 |
| 6h_C | 6h | 6 | 788598 | 34611 | 75437 | 823209 | 713161 | 0,0957 |
| 12h_A | 12h | 12 | 596322 | 54244 | 180882 | 650566 | 415440 | 0,3033 |
| 12h_B | 12h | 12 | 603069 | 49660 | 135560 | 652729 | 467509 | 0,2248 |
| 12h_C | 12h | 12 | 416693 | 23923 | 93748 | 440616 | 322945 | 0,225 |
| 24h_A | 24h | 24 | 371195 | 29709 | 141024 | 400904 | 230171 | 0,3799 |
| 24h_B | 24h | 24 | 441167 | 36707 | 171988 | 477874 | 269179 | 0,3898 |
| 24h_C | 24h | 24 | 381157 | 29220 | 137664 | 410377 | 243493 | 0,3612 |
| dRNA | MIX | NA | 895424 | 16925 | 318802 | 912349 | 576622 | 0,356 |
|  |  |  | 15062290 | 1087883 | 1475053 | 16150173 | 12831753 | 2,6012 |

The dRNA sequencing yielded 912,349 QC passed reads, out of which 318,802 was of viral origin, corresponding to a 35.6% of viral read ratio and a mean coverage of 244 across the viral genome (**Figure 4**). The two sequencing libraries compromise a total of 1,793,855 and 13,408,375 good quality viral and host reads, respectively.

Data (*bam* files containing the alignment and the sequence and its quality information as well) were uploaded to the European Nucleotide Archive, under the following BioProject: ***PRJEB56841***. All data can be used without restrictions.

### Technical Validation

#### RNA

Qubit RNA BR and HS Assay Kits (Invitrogen) were used to measure the amount of total RNA and polyA-selected RNA samples, respectively. The final concentrations of the RNA samples were determined by Qubit 4.0.

#### cDNA

The amount of the cDNA samples and the ready cDNA libraries were measured using Qubit 4.0 fluorometer and Qubit dsDNA HS Assay Kit (Invitrogen). The quality of RNA was detected with the Agilent 4150 TapeStation System. RNA samples with RIN values ≥ 9.0 were used for sequencing (**Figure 6**).

**Figure 6.**
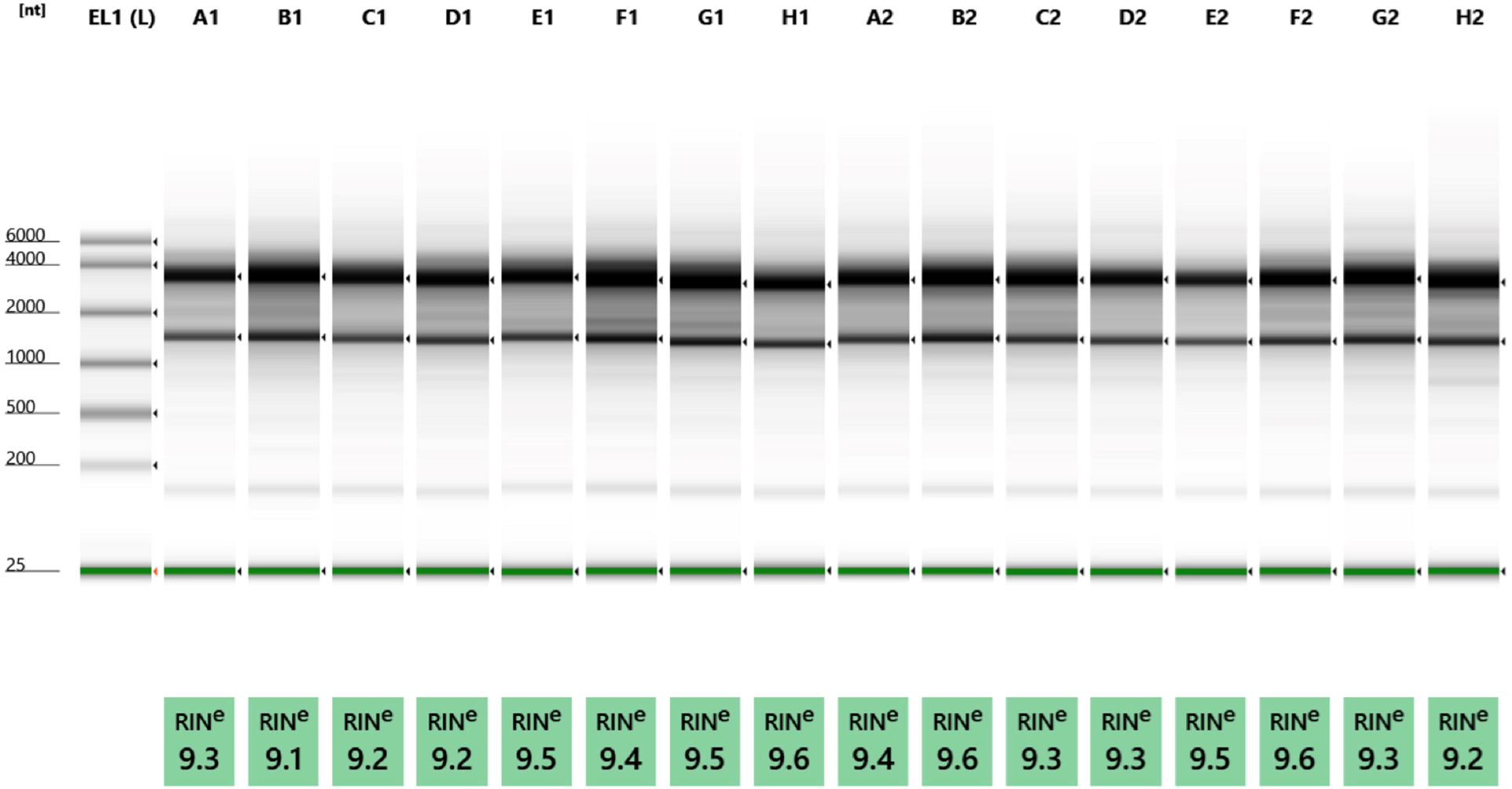
Quality of total RNA samples. The quality of the RNAs were assessed by using a TapeStation 4150 System and RNA ScreenTape (both from Agilent Technologies). TapeStation gel image shows that intact, high-quality RNAs (RIN > 9) were isolated from the cells and used for Nanopore sequencing. The image shows the following samples: EL1(L): marker; A1: 1h (replicate); B1: 1h (replicate C); C1: 2h (replicate A); D1: 2h (replicate B); E1: 4h (replicate A); F1: 4h (replicate B); G1: 6h (replicate A); H1: 6h (replicate B); A2: 12h (replicate A); B2: 12h (replicate C); C2: 24h (replicate A); D2: 24h (replicate B); E2: 2h (used for dRNA-seq); F2: 6h (used for dRNA-seq); G2: 12h (for dRNA-seq); H2: 24h (for dRNA-seq)

Three biological replicates were used for each of the infection time points. To analyze the effect of MPXV infection on the transcriptome profile of the host cells, mock-infected CV-1 cells were also harvested and sequenced.

### Usage Notes

Our dataset can be used to annotate novel viral transcripts and transcript isoforms, but possibly from the host as well. There are several bioinformatic tools that can be used to achieve this, including: TALON^45^; LIQA^46^; LoRTIA (https://github.com/zsolt-balazs/LoRTIA); EPI2ME’s transcriptomes workflow (https://github.com/epi2me-labs/wf-transcriptomes) or SQUANTI3 (https://github.com/ConesaLab/SQANTI3,^47^). Transcript annotation can be carried out from both types of sequencing data (dcDNA and dRNA), however as dRNA-seq yields less artificial or false products, it is suggested to use these reads for validating the dcDNA-seq derived transcripts^31^. Although it is possible that some rare transcripts that are expressed in a subset of the time-points exclusively (e.g., some immediate early isoforms) could not be captured in the dRNA sequencing library. After identification, the novel transcripts should be annotated to ORFs, their coding capacity be estimated, their TSS and TES sites be analyzed and accordingly their isoform categories be assessed (long or short TSS, alternative termination, etc.).

The gene-wise and/or transcript-wise gene counts from the cDNA-seq data can be subjected to differential gene expression (DGE) or differential transcript expression (DTE), respectively. Furthermore, differential transcript usage analyses (DTU) can be carried out as well, for example with RATS^48^. The https://github.com/nanoporetech/pipeline-transcriptome-depipeline, based loosely on the workflow presented in^49^, carries out these analyses from the annotated transcriptome, while EPI2ME’s transcriptomes workflow (https://github.com/epi2me-labs/wf-transcriptomes) carries out the transcript annotation and the above analyses in succession. The DGE, DTE and DTU analyses can be carried out both on the viral and on the host data and they can be based upon several comparisons, for example mock vs each time-point. In addition, the longitudinal expression data from cDNA-seq can be subjected to a time-series analysis as well^50^.

Besides focusing on individual genes or transcripts, gene-set enrichment analysis (GSEA) or pathway enrichment analyses can also be carried out to identify biological pathways that are affected by the viral infection in the host cells, for example with *pathfindR*^51^.

A combined workflow would be: 1.) detect transcripts using both sequencing approaches, but 2.) use the dRNA reads for validation, 3.) annotate them and carry out the transcript isoform analyses, 4.) quantify these validated transcripts in the cDNA data to estimate transcript counts, and finally 4.) carry out the above mentioned DGE, DTE, DTU and biological pathway analyses. Taken together, the almost 1.5 million viral and almost 13 million host reads enable the in-depth and temporal characterization of the Monkeypox transcriptome and the effect of the viral infection on the host’ gene expression.

## Code Availability

The complete workflow, from mapping to the generation of figures is available at the GitHub repository (https://github.com/Balays/MPOX_ONT_RNASeq).

## Acknowledgements

This research was supported by National Research, Development and Innovation Office (NRDIO), Researcher-initiated research projects (Grant numbers: K 128247 and K 142674) to ZB and by the NRDIO Research projects initiated by young researchers (Grant number: FK 128252) to DT. The work was also supported by National Laboratory of Virology (RRF-2.3.1-21-2022-00010) to GK and FJ. IP was supported by the New National Excellence Program of the Ministry for Innovation and Technology (ÚNKP-22-4-SZTE-310). ÁH was supported by Hungarian Ministry of Innovation and Technology, National Academy of Scientist Education, (FEIF/646–4/2021-ITM_SZERZ). The APC fee was covered by the University of Szeged, Open Access Fund: 5954.

## Author contributions

BK: carried out bioinformatics, analysis and interpretation of the data, wrote the manuscript

ÁD: took part in RNA isolation, carried out PolyA-selection and direct cDNA sequencing

ZC: isolated the total RNA and participated in dRNA sequencing

GK: carried out viral infection

JH: propagated the virus

DR: propagated the virus

IP: participated in data analysis and wrote the manuscript

VÉD: participated and interpretation of data

BD: propagated and maintained the CV-1 cell line

GT: participated in interpretation of data

FJ: supervision, participated in viral infection

GET: propagated the cells and the virus, participated in viral infection

FVF: propagated the cells and the virus, participated in viral infection

BZ: propagated the cells and the virus, participated in viral infection

ZL: propagated the cells and the virus, participated in viral infection

ÁH: participated in bioinformatics analysis

ÁF: participated in bioinformatics analysis

GG: participated in bioinformatics analysis

MM: participated in cell culture experiments

AAK: participated in bioinformatics analysis

DT: participated in the design of the experiments, in data analysis and wrote the manuscript

ZB: conceived and designed the experiments, supervised the project and wrote the manuscript.

All authors read and approved the final paper.

---

**Supplementary Figure 1.** Viral read length density distributions. **A)** Density plot for each sample with mean values shown as red dashed lines; **B)** Histogram for the combined cDNA and dRNA libraries; and **C)** Violin plot with added boxplots for each sample.

## References

1. Diven, D. G. An overview of poxviruses. J. Am. Acad. Dermatol. 44, 1–16 (2001).

2. Moss, B. & Smith, G. L. Poxviridae: The Viruses and Their Replication. in Field’s Virology 573–613 (2021).

3. Elwood, J. M. Smallpox and its eradication. J. Epidemiol. Community Heal. 43, 92–92 (1989).

4. Adler, H. et al. Clinical features and management of human monkeypox: a retrospective observational study in the UK. Lancet Infect. Dis. 22, 1153–1162 (2022).

5. Noe, S. et al. Clinical and virological features of first human monkeypox cases in Germany. Infection (2022). doi:10.1007/s15010-022-01874-z

6. Kumar, N., Acharya, A., Gendelman, H. E. & Byrareddy, S. N. The 2022 outbreak and the pathobiology of the monkeypox virus. J. Autoimmun. 131, 102855 (2022).

7. Luna, N. et al. Phylogenomic analysis of the monkeypox virus (MPXV) 2022 outbreak: Emergence of a novel viral lineage? Travel Med. Infect. Dis. 49, 102402 (2022).

8. Isidro, J. et al. Phylogenomic characterization and signs of microevolution in the 2022 multi-country outbreak of monkeypox virus. Nat. Med. 28, 1569–1572 (2022).

9. Kim, J. A., Park, S. K., Kumar, M., Lee, C. H. & Shin, O. S. Insights into the role of immunosenescence during varicella zoster virus infection (shingles) in the aging cell model. Oncotarget 6, 35324–35343 (2015).

10. Hendrickson, R. C., Wang, C., Hatcher, E. L. & Lefkowitz, E. J. Orthopoxvirus genome evolution: The role of gene loss. Viruses 2, 1933–1967 (2010).

11. Walsh, D. Poxviruses: Slipping and sliding through transcription and translation. PLoS Pathogens 13, (2017).

12. Alkhalil, A. et al. Gene expression profiling of monkeypox virus-infected cells reveals novel interfaces for host-virus interactions. Virol. J. 7, (2010).

13. Rubins, K. H., Hensley, L. E., Relman, D. A. & Brown, P. O. Stunned silence: Gene expression programs in human cells infected with monkeypox or vaccinia virus. PLoS One 6, (2011).

14. Bourquain, D., Dabrowski, P. W. & Nitsche, A. Comparison of host cell gene expression in cowpox, monkeypox or vaccinia virus-infected cells reveals virus-specific regulation of immune response genes. Virol. J. 10, (2013).

15. Rubins, K. H. et al. Comparative analysis of viral gene expression programs during poxvirus infection: A transcriptional map of the vaccinia and monkeypox genomes. PLoS One 3, 1–12 (2008).

16. Xuan, D. T. M. et al. Comparison of Transcriptomic Signatures between Monkeypox-Infected Monkey and Human Cell Lines. J. Immunol. Res. 2022, (2022).

17. Tombácz, D. et al. Time-course transcriptome profiling of a poxvirus using long-read full-length assay. Pathogens 10, 1–17 (2021).

18. Nagalakshmi, U., Waern, K. & Snyder, M. RNA-seq: A method for comprehensive transcriptome analysis. Current Protocols in Molecular Biology (2010). doi:10.1002/0471142727.mb0411s89

19. Mutz, K.-O., Heilkenbrinker, A., Lönne, M., Walter, J.-G. & Stahl, F. Transcriptome analysis using next-generation sequencing. Curr. Opin. Biotechnol. 24, 22–30 (2013).

20. Anamika, K., Verma, S., Jere, A. & Desai, A. Transcriptomic Profiling Using Next Generation Sequencing - Advances, Advantages, and Challenges. in Next Generation Sequencing: Advances, Applications and Challenges (ed. Kulski, J. K.) (IntechOpen, 2016). doi:10.5772/61789

21. Patterson, J. et al. Impact of sequencing depth and technology on de novo RNA-Seq assembly. BMC Genomics 20, 604 (2019).

22. Grünberger, F., Ferreira-Cerca, S. & Grohmann, D. Nanopore sequencing of RNA and cDNA molecules in Escherichia coli. RNA 28, 400–417 (2022).

23. Torma, G. et al. Combined Short and Long-Read Sequencing Reveals a Complex Transcriptomic Architecture of African Swine Fever Virus. Viruses 13, 579 (2021).

24. Torma, G. et al. Dual isoform sequencing reveals complex transcriptomic and epitranscriptomic landscapes of a prototype baculovirus. Sci. Rep. 12, 1291 (2022).

25. Shchelkunov, S. N. et al. Analysis of the Monkeypox Virus Genome. Virology 297, 172–194 (2002).

26. Prazsák, I. et al. Long-read sequencing uncovers a complex transcriptome topology in varicella zoster virus. BMC Genomics 19, 1–20 (2018).

27. Soneson, C. et al. A comprehensive examination of Nanopore native RNA sequencing for characterization of complex transcriptomes. Nat. Commun. 10, 1–14 (2019).

28. Depledge, D. P. et al. Direct RNA sequencing on nanopore arrays redefines the transcriptional complexity of a viral pathogen. Nat. Commun. 10, 1–13 (2019).

29. Olasz, F. et al. Short and Long-Read Sequencing Survey of the Dynamic Transcriptomes of African Swine Fever Virus and the Host Cells. Front. Genet. 11, (2020).

30. Fülöp, Á. et al. Integrative profiling of Epstein–Barr virus transcriptome using a multiplatform approach. Virol. J. 19, 7 (2022).

31. Tombácz, D. et al. In-Depth Temporal Transcriptome Profiling of an Alphaherpesvirus Using Nanopore Sequencing. Viruses 14, 1289 (2022).

32. Yang, Z., Bruno, D. P., Martens, C. A., Porcella, S. F. & Moss, B. Simultaneous high-resolution analysis of vaccinia virus and host cell transcriptomes by deep RNA sequencing. Proc. Natl. Acad. Sci. U. S. A. (2010). doi:10.1073/pnas.1006594107

33. Yang, Z., Bruno, D. P., Martens, C. A., Porcella, S. F. & Moss, B. Genome-Wide Analysis of the 5’ and 3’ Ends of Vaccinia Virus Early mRNAs Delineates Regulatory Sequences of Annotated and Anomalous Transcripts. J. Virol. (2011). doi:10.1128/jvi.00428-11

34. Yang, Z. et al. Expression Profiling of the Intermediate and Late Stages of Poxvirus Replication. J. Virol. (2011). doi:10.1128/jvi.05446-11

35. Yang, Z., Martens, C. A., Bruno, D. P., Porcella, S. F. & Moss, B. Pervasive initiation and 3’-end formation of poxvirus postreplicative RNAs. J. Biol. Chem. (2012). doi:10.1074/jbc.M112.390054

36. Yang, Z., Maruri-Avidal, L., Sisler, J., Stuart, C. A. & Moss, B. Cascade regulation of vaccinia virus gene expression is modulated by multistage promoters. Virology (2013). doi:10.1016/j.virol.2013.09.007

37. Tombácz, D. et al. Long-read assays shed new light on the transcriptome complexity of a viral pathogen. Sci. Rep. 10, 1–13 (2020).

38. Tombácz, D. et al. Dynamic transcriptome profiling dataset of vaccinia virus obtained from long-read sequencing techniques. Gigascience 7, (2018).

39. Maróti, Z. et al. Time-course transcriptome analysis of host cell response to poxvirus infection using a dual long-read sequencing approach. BMC Res. Notes 14, 239 (2021).

40. Li, H. Minimap2: Pairwise alignment for nucleotide sequences. Bioinformatics (2018). doi:10.1093/bioinformatics/bty191

41. Wickham, H. et al. Welcome to the Tidyverse. J. Open Source Softw. 4, 1686 (2019).

42. Morgan M, Pagès H, Obenchain V, H. N. Rsamtools: Binary alignment (BAM), FASTA, variant call (BCF), and tabix file import. (2022).

43. Wickham, H. ggplot2. (Springer New York, 2009). doi:10.1007/978-0-387-98141-3

44. Gu, Z., Gu, L., Eils, R., Schlesner, M. & Brors, B. circlize implements and enhances circular visualization in R. Bioinformatics 30, 2811–2812 (2014).

45. Wyman, D. et al. A technology-agnostic long-read analysis pipeline for transcriptome discovery and quantification. bioRxiv 672931 (2020). doi:10.1101/672931

46. Hu, Y. et al. LIQA: long-read isoform quantification and analysis. Genome Biol. (2021). doi:10.1186/s13059-021-02399-8

47. Tardaguila, M. et al. SQANTI: Extensive characterization of long-read transcript sequences for quality control in full-length transcriptome identification and quantification. Genome Res. (2018). doi:10.1101/gr.222976.117

48. Froussios, K., Mourão, K., Simpson, G., Barton, G. & Schurch, N. Relative abundance of transcripts (RATs): Identifying differential isoform abundance from RNA-seq [version 1; referees: 1 approved, 2 approved with reservations]. F1000Research 8, 1–21 (2019).

49. Love, M. I., Soneson, C. & Patro, R. Swimming downstream: Statistical analysis of differential transcript usage following Salmon quantification. F1000Research (2018). doi:10.12688/f1000research.15398.3

50. Varoquaux, N. & Purdom, E. A pipeline to analyse time-course gene expression data. F1000Research 9, 1447 (2020).

51. Ulgen, E., Ozisik, O. & Sezerman, O. U. PathfindR: An R package for comprehensive identification of enriched pathways in omics data through active subnetworks. Front. Genet. (2019). doi:10.3389/fgene.2019.00858

