## Supplementary figures and images for "In-depth Temporal Transcriptome Profiling of Monkeypox and Host Cells using Nanopore Sequencing"

### Supplementary Figure 1

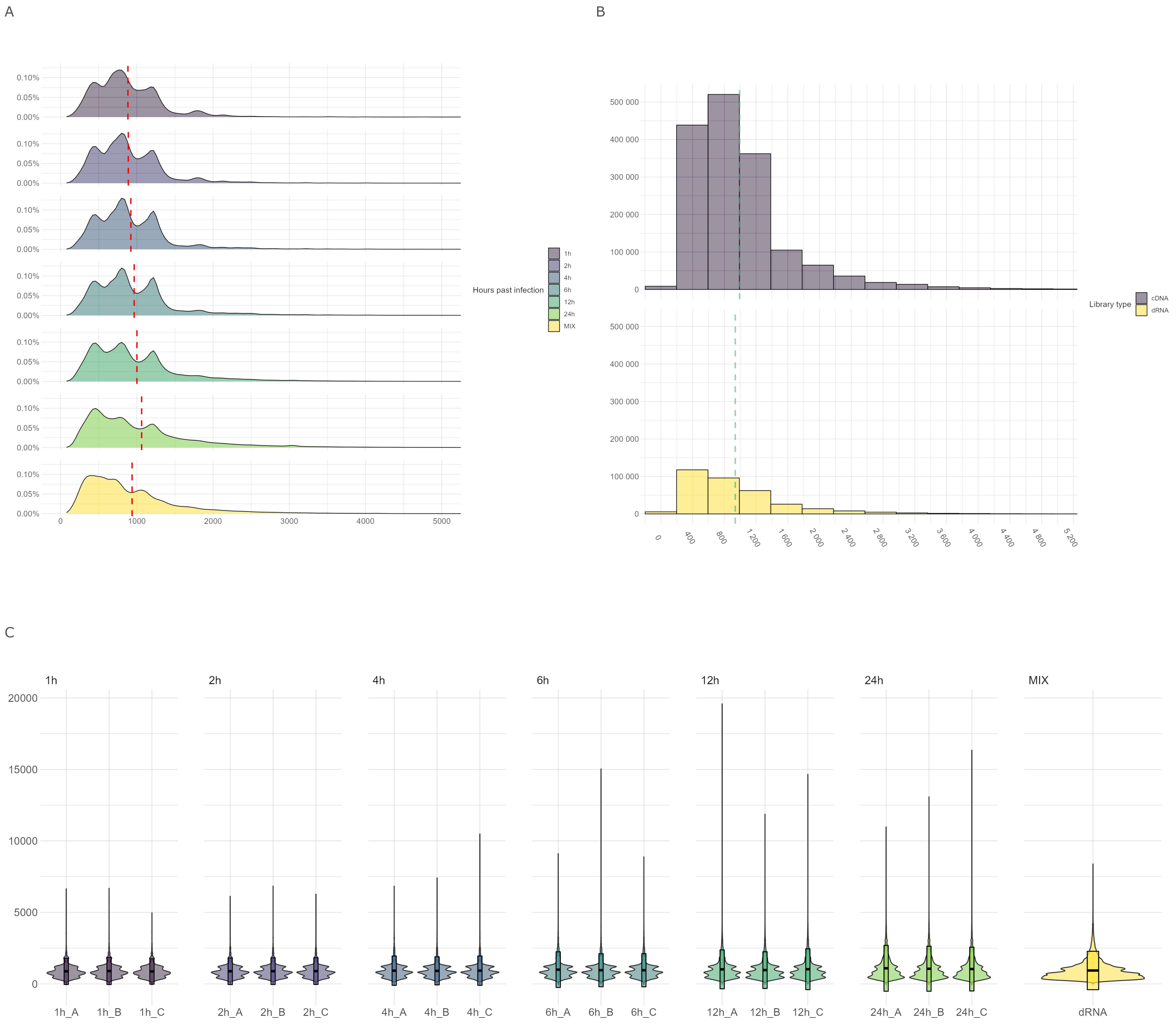
